# Intra-tumor heterogeneity of Diffuse Large B-cell Lymphoma involves the induction of diversified stroma-tumor interfaces

**DOI:** 10.1101/2020.05.30.124859

**Authors:** Sabina Sangaletti, Fabio Iannelli, Federica Zanardi, Valeria Cancila, Paola Portararo, Laura Botti, Davide Vacca, Claudia Chiodoni, Arianna Di Napoli, Cesare Valenti, Maria Carmela Vegliante, Federica Pisati, Alessandro Gulino, Maurilio Ponzoni, Mario Paolo Colombo, Claudio Tripodo

## Abstract

Intra-tumor heterogeneity in lymphoid malignancies is articulated around several fundamentals, encompassing selection of genetic subclonal events and epigenetic regulation of transcriptional programs. Clonally-related neoplastic cell populations are unsteadily subjected to immune editing and metabolic adaptations within different tissue microenvironments. How tissue-intrinsic mesenchymal determinants impact on the diversification of aggressive lymphomas is still unknown. In this study we adopted the established A20 line-based model of Diffuse Large B-cell Lymphoma (DLBCL), to investigate the intra-tumor heterogeneity associated with the infiltration of different tissue microenvironments and the specific mesenchymal modifications that characterize stromal adaptation to lymphoma seeding. Combining *in situ* quantitative immunophenotypical analyses and RNA sequencing, we found that the tissue microenvironment casts a relevant influence over A20 transcriptional landscape also impacting on Myc and DNA damage response programs. Extending the investigation to mice deficient for the matricellular protein Sparc, a stromal determinant endowed with a strong prognostic significance in human DLBCL, we demonstrated a different immune imprint on A20 cells related to stromal *Sparc* proficiency. The study provides the first evidence of human DLBCL intra-lesional heterogeneity arising from diversified mesenchymal contextures and impacting on MYC expression, advancing the challenge for resolving intra-tumor heterogeneity probing stromal/immune interfaces.

**KEY POINTS:**

1. Diversified stromal adaptations of infiltrated tissues shape DLBCL intra-tumor heterogeneity regulating transcriptional/phenotypic features
2. Stromal Sparc, which endorses prognostic significance in DLBCL, tunes the immune pressure exerted on lymphoma cells by the microenvironment

## INTRODUCTION

In the era of liquid biopsy-based analyses of the systemic tumor landscape of patients, the exploitation of events on a subclonal scale has enabled a new level of management of intra-tumor heterogeneity in lymphomas.^1^ If the systemic overview of clonal evolution offers new strategies for prognostication and new windows of opportunity on targeted treatments,^2^ the nature of local dynamics contributing to such diversification are still poorly defined and the role of tissue stromal and immune microenvironments in influencing clonal diversity at phenotypic, transcriptional, or genetic levels deserves investigation.

Diffuse Large B-cell Lymphoma (DLBCL) represents a highly heterogeneous diagnostic category including aggressive malignant proliferations of mature B cells with variable imprints of germinal center (GC)-derivation or post-GC differentiation - referred to as cell of origin (COO) - and dramatic variability in mutational signatures.^3^ The most recent efforts in the interpretation and classification of DLBCL heterogeneity have relied on the integration of genetic and transcriptional signatures and led to the identification of discrete clusters characterized by different oncogenic trajectories, biologies, and clinical behavior.^4-5^ In the attempt to deconvolve the DLBCL complexity, the nature of associated stromal microenvironment has been investigated, revealing several layers of complexity in the stromal microenvironment, some of which associated with the COO, others providing further diversification.^6-7^ From the seminal studies by the Staudt’s group, a prognostic significance of the DLBCL-associated stromal microenvironment clearly emerged, which identified among the major determinants the matricellular protein SPARC, by the authors associated with tumor-infiltrating macrophages.^6,8^ Indeed, *SPARC* consistently emerged as the fronting gene of prognostically relevant microenvironment-related signatures in DLBCL,^6-7,9^ being low *SPARC* levels associated with poor prognosis in immuno-chemotherapy-treated DLBCL patients. Our group has previously demonstrated that Sparc is a major stromal factor supporting bone marrow (BM) B lymphopoiesis and secondary lymphoid organ (SLO) function influencing the mesenchymal architecture of the GC.^10-11^ Moreover, we demonstrated that defective Sparc expression in SLO licensed the activation of myeloid elements towards class-I Interferon-driven responses eventually unleashing malignant lymphoproliferation in the setting of persistent immune stimulation.^12^

In the present study we investigated whether tissue-specific microenvironmental cues may influence the phenotypic and transcriptional heterogeneity of an established full-blown aggressive clonal DLBCL model based on the A20 cell line.^13^ We focused on two levels of microenvironment complexity: one related to the multiple tissue localizations of the lymphomatous cells, namely bone marrow (BM), liver (LI), and spleen (SPL); the other related to Sparc proficiency/deficiency in the stroma. We demonstrated that the A20 transcriptome was substantially modulated by the different host tissue microenvironments, which gave rise to diversified stromal modifications upon A20 infiltration. We found that Myc expression and its related transcriptional programs - along with DNA damage pathways - were affected by the tissue environment. We observed that Sparc-defective stroma induced the overexpression of Complement-associated innate immune/inflammatory programs and affected the expression of oncogenic/metabolic pathways thus exerting a higher immune pressure on lymphomatous cells. Moreover, we report the presence of mesenchymal intra-lesional diversity matching MYC spatial heterogeneity in human DLBCLs.

## METHODS

### Mice and cell lines

BALB/cAnNCrl mice were purchased from Charles River Laboratories (Calco, Italy). *Sparc*^*-/-*^ mice (on a BALB/c background) were generated and maintained in the Molecular Immunology Unit of the National Cancer Institute, Milan Italy.^14^ All the experiments involving animals described in this study were approved by the Ministry of Health (authorization number 1027/2016-PR). BALB/c-derived A20 B lymphoma cell line was obtained from the American Type Culture Collection (Rockville, MD) and maintained in RPMI 1640 supplemented with 10% FCS (GIBCO). For in vivo experiments A20 cell lines were injected i.v. (5*10^5^ cells). To perform multiple analyses and easily follow A20 lymphomatous cells, A20-GFP+ cells were adopted in multiple transplantation experiments (see Supplemental Methods).

### Human DLBCL tissue samples

Formalin-fixed and paraffin embedded (FFPE) tissue samples of human DLBCL cases diagnosed between 2017 and 2019 at the San Raffaele Hospital Haematopathology Unit, Milan, Italy and at the Sant’Andrea “La Sapienza” University Hospital, Rome, Italy, were selected for the quantitative *in situ* immunophenotypical and multiresolution analyses. The cases were selected according to a variable expression of MYC - ranging from 10% to 60% and characterized by zonal/focal distribution. The clinical-pathological characteristics of the 12 cases are summarized in Supplemental Table S1. Samples were collected according to the Helsinki Declaration and the study was approved by the University of Palermo Institutional Review Board (approval number 09/2018).

### In situ histomorphological and quantitative immunophenotypical analyses

Tissues dissected from mice were washed in PBS and collected for fixation in 10% neutral buffered formalin overnight, washed in water and paraffin-embedded. Bone samples were decalcified using an EDTA-based decalcifying solution (MicroDec EDTA-based, Diapath) for 8 hours, then washed out with water for 1 hour and subsequently processed and embedded in paraffin.

Four-micrometers-thick sections from mouse FFPE tissues were stained for H&E to define tissue infiltration and tumor morphology. Immunohistochemical and immunofluorescence stainings were performed as previously reported^12^ and detailed in the Supplemental Methods section. For multiple-marker immunostainings, sections were subjected to sequential rounds of single-marker immunostaining and the binding of the primary antibodies was revealed by the use of specific secondary antibodies conjugated with different fluorophores or enzymes (See Supplemental Methods section). Slides were analyzed under a Zeiss Axioscope A1 microscope equipped with four fluorescence channels widefield IF. Microphotographs were collected using a Zeiss Axiocam 503 Color digital camera with the Zen 2.0 Software (Zeiss). Slide digitalization was performed using an Aperio CS2 digital scanner (Leica Biosystems) with the ImageScope software.

Quantitative analyses of immunohistochemical stainings were performed by calculating the average percentage of positive signals in five non-overlapping fields at high-power magnification (x400) using the Nuclear Hub or Positive Pixel Count v9 Leica Software Image Analysis.

Quantitative analyses of immunofluorescence stainings were performed in five non-overlapping fields at high-power magnification (x400), by isolation of the fluorescent marker component and measurement of its density. Multi-resolution analysis of double-marker immunostained sections has been performed through an ad-hoc developed software tool (See Supplemental Methods).^15-16^

### Flow cytometry

To evaluate A20 lymphoma take in WT and *Sparc*^*-/-*^ mice in BM, LI, and SPL, and to recover A20 cells from the BM such to perform re-transplantation experiments, cell suspensions were stained with antibodies to CD45, CD19 and B220. A20 cells can be discriminated from the normal B-cell counterpart according to their higher expression of CD45 and B220 combined with a higher side scatter (SSC-A) (Supplemental Figure S1). To characterize T cell infiltrates in WT and *Sparc*^*-/*-^ mice in re-transplantation experiments, BM cell suspensions were stained with antibodies to CD4, CD8, Foxp3, Ki-67, TIM3, ICOS, CD25, PD1, OX40, H-2Kd, I-A/I-E. The antibodies used are detailed in the Supplemental Methods section. Surface staining was performed in phosphate-buffered saline (PBS) supplemented with 2% fetal bovine serum (FBS) for 30 min on ice. Foxp3 intracellular staining was performed according to the manufacturer’s instructions (eBioscience). Flow cytometry data were acquired on a LSRFortessa (Becton Dickinson) and analyzed with FlowJo software (version 8.8.6, Tree Star Inc.).

### Mesenchymal cell purification and in vitro co-culture experiments

Murine BM mesenchymal stromal cell (MSC) cultures were obtained from the trabecular fraction of femurs and tibias of WT and *Sparc*^*-/-*^ mice. Briefly, the cellular fraction of the femurs and tibias was washed out and the compact bone was incubated with collagenase I (1 mg/ml) for 1 h at 37°C. After enzyme digestion, the bone suspension was passed through a 70μm filter mesh to remove debries. Cells were seeded in complete medium (MesenCult Basal Medium) at a density of 25*10^6^ cells/ml. Floating cells were removed every 3-4 days. Adherent cells were phenotypically characterized using the following antibodies: CD31, CD45, CD34, cKit, Ter119, CD44, Sca-1, CD29. The antibodies adopted are detailed in the Supplemental Methods Section. BM-MSCs were defined according to their negativity for hematopoietic and lineage markers (CD45, CD34, CD31, cKit, Ter119) and positivity for the MSC-markers CD44, CD29 and Sca-1. Perycytic mesenchymal cells (An2+, the mouse homolog of NG2) were enriched from BM cell suspension using anti-An2/NG2 microbeads from Miltenyi. In vitro experiments involving murine An2+MSCs were performed using cells between the 2nd and the 5th passages. For co-culture experiments 10^5^ BM-MSCs or An2+MSCs were seeded over-night into a 24-well plate and then added with 10^5^ A20 cells. At basal, 48h, and 96h time points, A20 cells were recovered and processed for qPCR analysis. To analyze Myc expression qPCR primers from TaqMan (Myc: Mm00487804_m1) were adopted.

### RNA sequencing and data analysis

Whole-transcriptome stranded sequencing libraries were generated using the TruSeq Stranded Total RNA with Ribo-Zero in order to remove ribosomal RNA. Sequencing was carried out on the Illumina HiSeq 3000 system requiring 80M reads/sample with paired-end mode. Reads were aligned to the ENSEMBL mouse genome assembly BALB_cJ_v1 using STAR.

Differentially expressed genes were called using DESeq2 R package starting from the count tables generated by HTSeq.

PCA plot was generated with the function plotPCA of the DESeq2 package, using default parameters on the log2 transformed intensity values. Euclidean distance between samples was calculated with the dist function of DESeq2 using default parameters. Gene fusion events were detected in RNA-seq data using the tool FusionCatcher (https://github.com/ndaniel/fusioncatcher). Additional details regarding the RNA sequencing data analysis are reported in the Supplemental Methods section.

### Gene set enrichment analysis, hierarchical clustering, and CIBERSORT deconvolution on human DLBCL cases

GSEA^17^ was performed on the GSE98588 dataset^4^ using the continuous expression level of NGFR gene to label the cases (ranked by Pearson metric method). We then tested roughly 300 gene sets extracted from widely used annotated databases (Kegg, Reactome, MSigDB) including several cellular processes and MYC-related pathways.

Raw data from Chapuy et al.^4^ were collected and used to generate expression profiles by RMAExpress (Robust Multi-Array Average). Multiple probes were collapsed to unique genes by selecting the ones with the maximal average expression for each gene. Expression values were log2 transformed. Heatmap and clustering analysis were performed using R statistical software. For hierarchical clustering analysis the Euclidian distances across samples were calculated and used to applied complete cluster aggregation method within pheatmap R package.

For CIBERSORT ^18^ analyses, we created a customized signature matrix as previously reported.^7^ Briefly, we used GEP from 24 different purified cell types collected from published datasets, to generate a microenvironment matrix specific for DLBCL.^7^ The customized signature featuring 990 genes characteristic of tumor samples of both ABC and GCB COO categories, immune cells and stromal compartment, was applied to the 137 DLBCLs from Chapuy et al.^4^ The 137 cases were divided into two groups according to the median value of *SPARC* transcript expression from quantile normalized and log2 transformed gene expression measurements.

### Statistical analysis

Unless differently specified, data are presented as the mean ± standard error of the mean. Two classes comparisons were performed according to either two-tailed Student’s T test or contingency two-sided Fisher’s exact test for discrete variables. For multiple classes comparisons, one-way analysis of variance (ANOVA) was performed. A p<0.05 was considered significant and indicated with *; p<0.01 was indicated as ** and p<0.001 as ***.

## RESULTS

### A20 DLBCL infiltrates display different morphology and phenotype depending on the tissue microenvironment

The tissue microenvironment-induced heterogeneity of A20 lymphomas was initially investigated by histomorphological and immunophenotypical characterization of tissue infiltrates in different sites upon i.v. injection of 5×10^5^ cells into BALB/c mice. Following 5 weeks of injection, mice were sacrificed and the A20 cells recovered from the BM, LI, and SPL tissues through flow cytometry sorting (Supplemental Figure S1) and processed for total RNA-sequencing as described in the Supplemental Methods section. BM, LI, and SPL from 5 other mice were collected for histopathology (Figure 1A). To specifically assess the relevance of stromal SPARC-induced modifications in A20 transcriptional programs, A20 cells were also injected into *Sparc*^*-/-*^ BALB/c hosts (Figure 1A).

**Figure 1.**
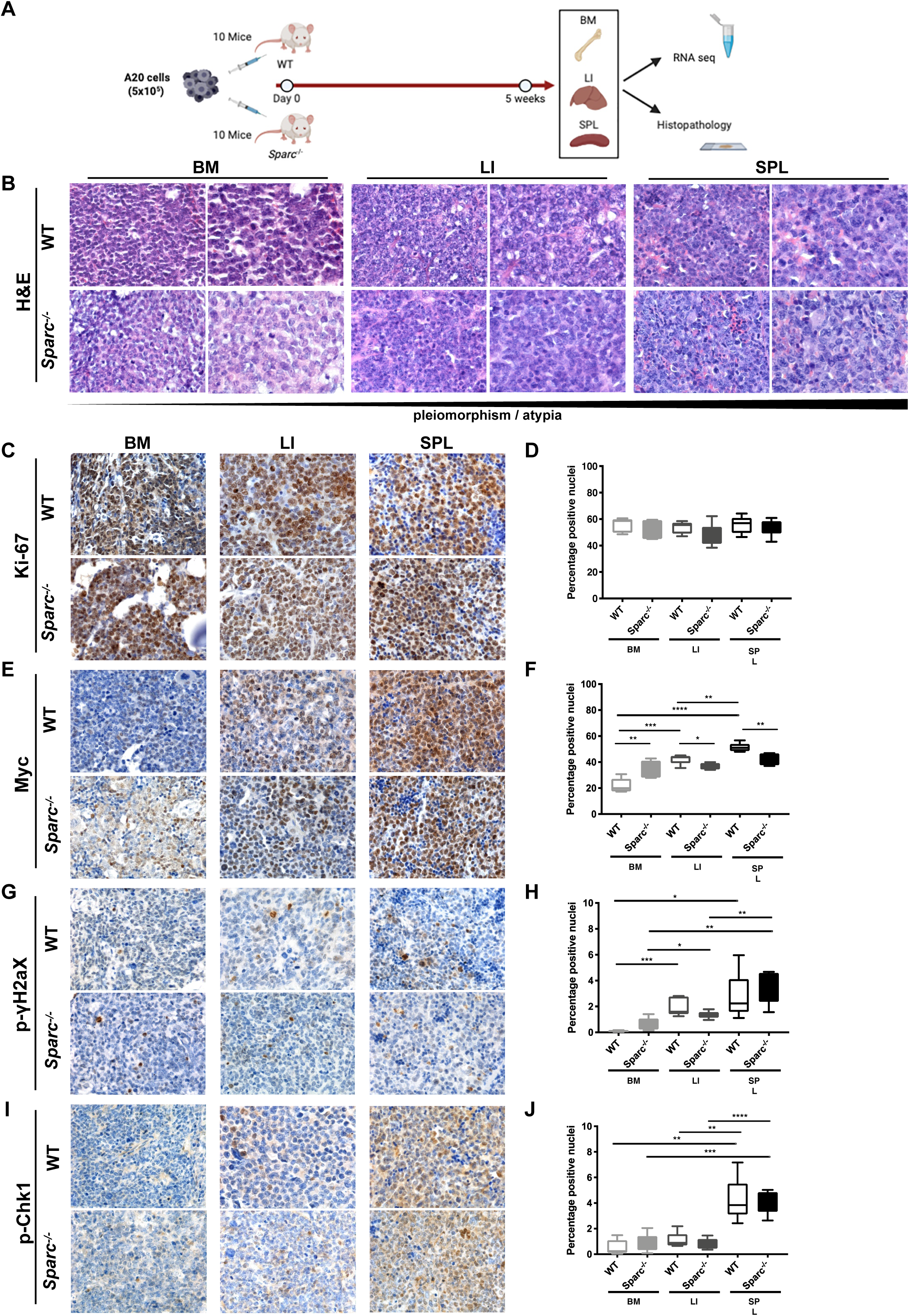
A20 DLBCL tumors display morphological and phenotypical heterogeneity in different tissue sites. (A) Graphical scheme of the in vivo A20 injection and purification experiment aimed at investigating the host tissue microenvironment effect on A20 morphology, phenotype and transcriptome. (B) Representative microphotographs of H&E-stained sections from A20 DLBCL infiltrates showing the increasing degree of pleiomorphism and cytological atypia of A20 cells from BM to LI and SPL tissues in WT and *Sparc*^*-/-*^ hosts. Original magnification x400 and x630. C-D, IHC for Ki-67 (C) and quantitative analysis (D) showing comparable proliferative fractions in BM, LI, and SPL tissue infiltrates. (E-F) IHC for Myc (E) and quantitative analysis (F) highlighting significant differences in Myc nuclear expression among BM, LI, and SPL A20 infiltrates. G-J IHC for p-gH2aX and p-Chk1 DNA damage response markers (G, I) and quantitative analysis (H, J) showing an increase fraction of immunoreactive nuclei in A20 SPL infiltrates as compared with LI and, even more significantly, BM. Original magnifications x400.

A20 cells formed diffuse infiltrates within the BM parenchyma, which mostly consisted of monomorphic medium-sized cells with hyperchromatic nuclei in WT and *Sparc*^*-/-*^ hosts (Figure 1B). In the LI, A20 cells infiltrated the hepatocyte muralia, forming multiple nodular foci mostly composed by atypical large cells with a high degree of pleiomorphism and scattered anaplastic figures (Figure 1B). The SPL architecture was almost completely effaced by the proliferation of A20 atypical cells infiltrating the red pulp and also extending to white pulp remnants, showing the highest degree of morphological atypia and frequency of mitotic figures with no relevant difference between the WT and *Sparc*^*-/-*^ genotypes (Figure 1B). The observed degree of morphological variation among BM LI and SPL in the A20 tumors was reminiscent of reports on human DLBCL with discordant morphology between lymphoid tissue and BM foci, with BM DLBCL infiltrates showing a lower degree of morphological atypia and smaller cytology.^19^ Moreover, it suggested that underlying differences in biological features of the malignant clone could be induced by tissue site-intrinsic adaptations.

To challenge this hypothesis, we assessed the proliferation of A20 cells in the different tissue sites, along with the expression of the key transcription factor Myc, which is known to drive tumor aggressiveness in B-cell lymphomas.^20^ Quantitative IHC analysis of the proliferative index of A20 DLBCL infiltrates assessed as the percentage of Ki-67+ cells revealed no significant differences among BM SPL and LI infiltrates in both the WT and *Sparc*^*-/-*^ settings (Figure 1C-D). Notably, A20 DLBCL infiltrates showed consistent variation in the extent of Myc expression, which was significantly higher in the SPL infiltrates than in LI and BM, with a higher trend in WT as compared with *Sparc*^*-/-*^ hosts, excepted for the BM where *Sparc*^*-/-*^ samples showed a higher expression than WT (Figure 1E-F). This finding indicates that despite a comparable proliferative attitude, A20 lymphomatous cells populating different tissue microenvironments could be induced to modulate key transcriptional regulators, such as Myc.

Myc expression facilitates the acquisition of a mutator phenotype in malignant B cells, by promoting DNA replicative stress and DNA damage induction. We therefore investigated the expression of DNA damage and replication stress markers phospho (p)-γH2ax and p-Chk1 by *in situ* quantitative IHC analysis. Consistently with Myc expression, p-γH2ax and p-Chk1 were significantly more expressed within SPL infiltrates as compared with LI and BM and showed a trend towards decrease in the *Sparc*^*-/-*^ genotype excepted for *Sparc*^*-/-*^ BM infiltrates, which showed slightly higher expression of the two markers (Figure 1 G-J). Collectively these data point to the selection of different DNA damage-prone phenotypes within different tissue contextures.

### Diversified mesenchymal adaptations characterize human DLBCL nodal lesions and A20 tissue infiltrates

On the basis of the observed morphological and phenotypic heterogeneity in the A20 lymphomas among different tissue sites, we investigated whether host tissue-related features could contribute to the diversification of the three microenvironments and/or of the two *Sparc* genotypes.

Human DLBCLs are associated with extremely diversified types of stromal adaptations, which have been put into relation with the cell of origin and phenotype of the malignant cells^.6-7^ thus configuring a high level of intra-tumor complexity.

In a small set of nodal DLBCL cases (n=12), comprising cases of GC and non-GC phenotype according to Hans algorithm^21^ and selected according to the heterogeneous, focal/zonal, pattern of MYC expression (Supplemental Table S1), we highlighted the presence of different types of mesenchymal cell meshworks within distinct areas of the same nodal lesions (Supplemental Figure S2A). In these cases, SMA+ myofibroblastic/reticular cells, NGFR+ (CD271) and CD146+ pericytic/mesenchymal stromal cells, and PDGFRβ+ perivascular stromal cells showed disconnected patterns (Supplemental Figure S2A). Moreover, mesenchymal markers including NGFR, SMA, PDGFRβ, and VCAM-1 that were either spatially associated or mutually exclusive in the GC or peri-follicular areas of non-neoplastic lymph nodes, were aberrantly splitted or focally associated in DLBCL infiltrates (Supplemental Figure S2B). Of note, within DLBCL nodal infiltrates, prominent heterogeneity in the density and spatial segregation of mesenchymal meshworks formed by different cell types was observed. According to NGFR and SMA double-marker immunostainings, DLBCL lesions showed dense, moderate or low NGFR and SMA mesenchymal meshworks either spatially segregated or nterweaved (Figure 2A).

**Figure 2.**
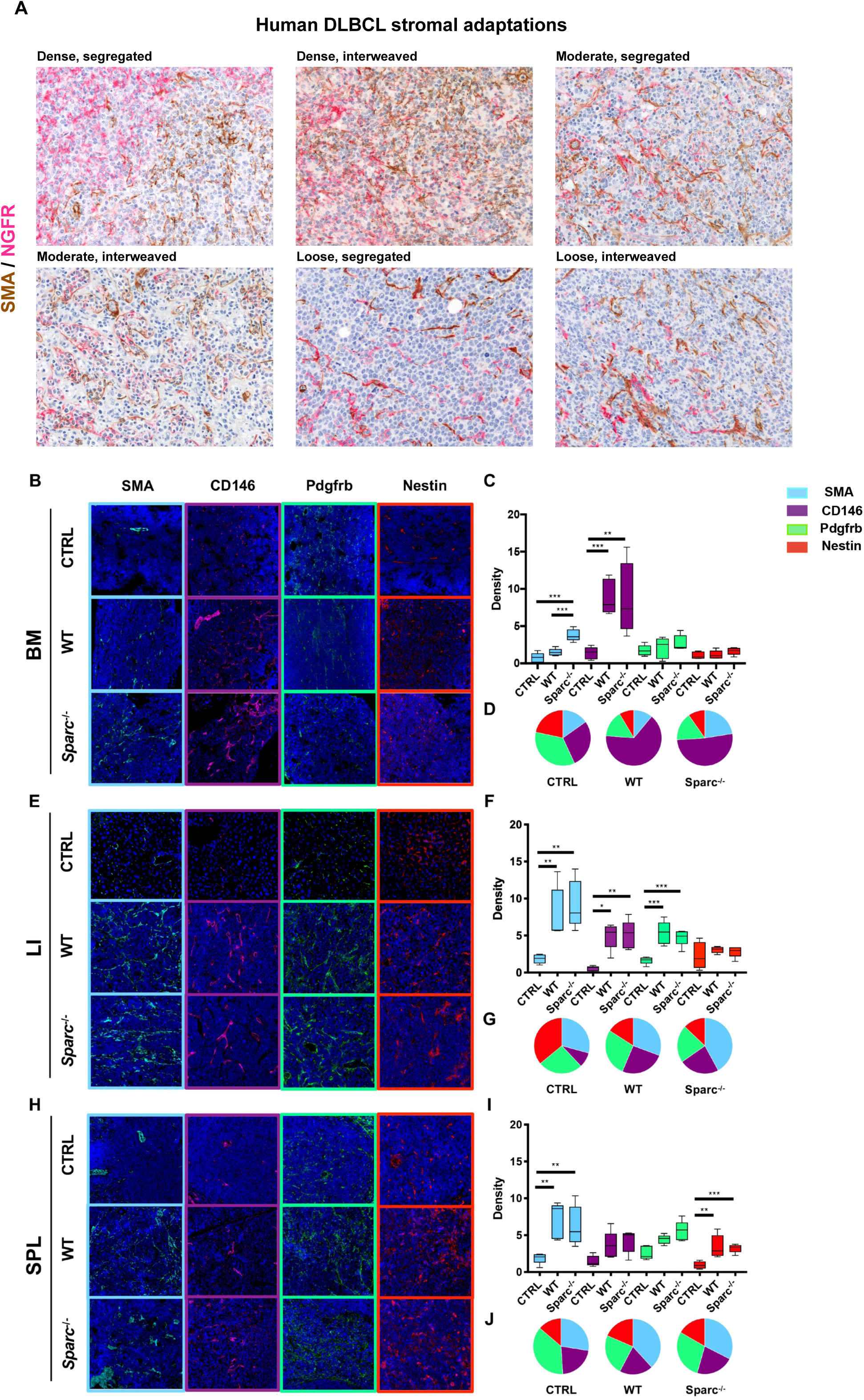
Mesenchymal meshwork heteogeneity in human DLBCL nodal lesions and in A20 infiltrates at different tissue sites. (A) Double-marker IHC for SMA (brown signal) and NGFR (purple signal) in the nodal infiltrates of six representative DLBCL cases (Supplemental Table S1) reflecting the variable degree (dense, moderate, low) of stromal reaction and spatial segregation (segregated, interweaved) of the two mesenchymal meshworks, supporting the engendering of diversified tumor-stroma interfaces. Original magnifications x200. (B-J) IF for SMA (cyan signal), CD146 (violet signal), Pdgfrb (green signal), and Nestin (red signal) mesenchymal markers (B, E, H), quantitative analysis (histograms, C,F,I) and average relative fractions (pie charts, D, G, J) of uninvolved (CTRL) or infiltrated BM (B-D), LI (E-G), and SPL (H-J) from WT and *Sparc*^*-/-*^ hosts, detailing that different combinations of mesenchymal markers are variably induced across the three tissue sites. Original magnifications x200.

Prompted by the evidence of an active and variably intense mesenchymal response to human DLBCL infiltration contributing to intra-tumor heterogeneity, we tested the hypothesis that host tissue adaptation to A20 lymphoma seeding could imply a diversified remodeling of mesenchymal cell meshworks. We thus analyzed the density of Sma, Cd146, Pdgfrb and Nestin mesenchymal markers in the BM, LI and SPL A20 infiltrates by quantitative immunofluorescence. In the BM, a prominent induction in Cd146+ elements characterized A20 lymphomas in both the WT and *Sparc*^*-/-*^ hosts, while Sma+ stromal cells were significantly induced only in *Sparc*^*-/-*^ infiltrates (Figure 2B-D). In LI, A20 tumor microenvironment was marked by a significant induction of Sma+, Cd146+, and Pdgfrb+ mesenchymal elements in both genotypes (Figure 2E-G), while the stromal reaction of SPL foci was characterized by a significant expansion of Sma+ and Nestin+ cells (Figure 2H-J). These results suggest that different qualities of stromal tissue adaptation to A20 DLBCL infiltration occur, which provide a microenvironment frame for the observed variations in the malignant clone phenotype.

### A20 lymphoma cells gene expression profile is influenced by the tissue microenvironment

We then analyzed by RNA-seq, the transcriptome of A20 cells purified from the three environments of WT and *Sparc*^*-/-*^ mice (Supplemental Table S2). Principal component analysis of the samples showed a predominance of tissue site over *Sparc* genotype in determining the clustering of the A20 lymphoma samples (Supplemental Figure S3A-B). Indeed, unsupervised clustering analysis of A20 DLBCL transcriptome identified clusters on the basis of the tissue environment (Figure 3A).

**Figure 3.**
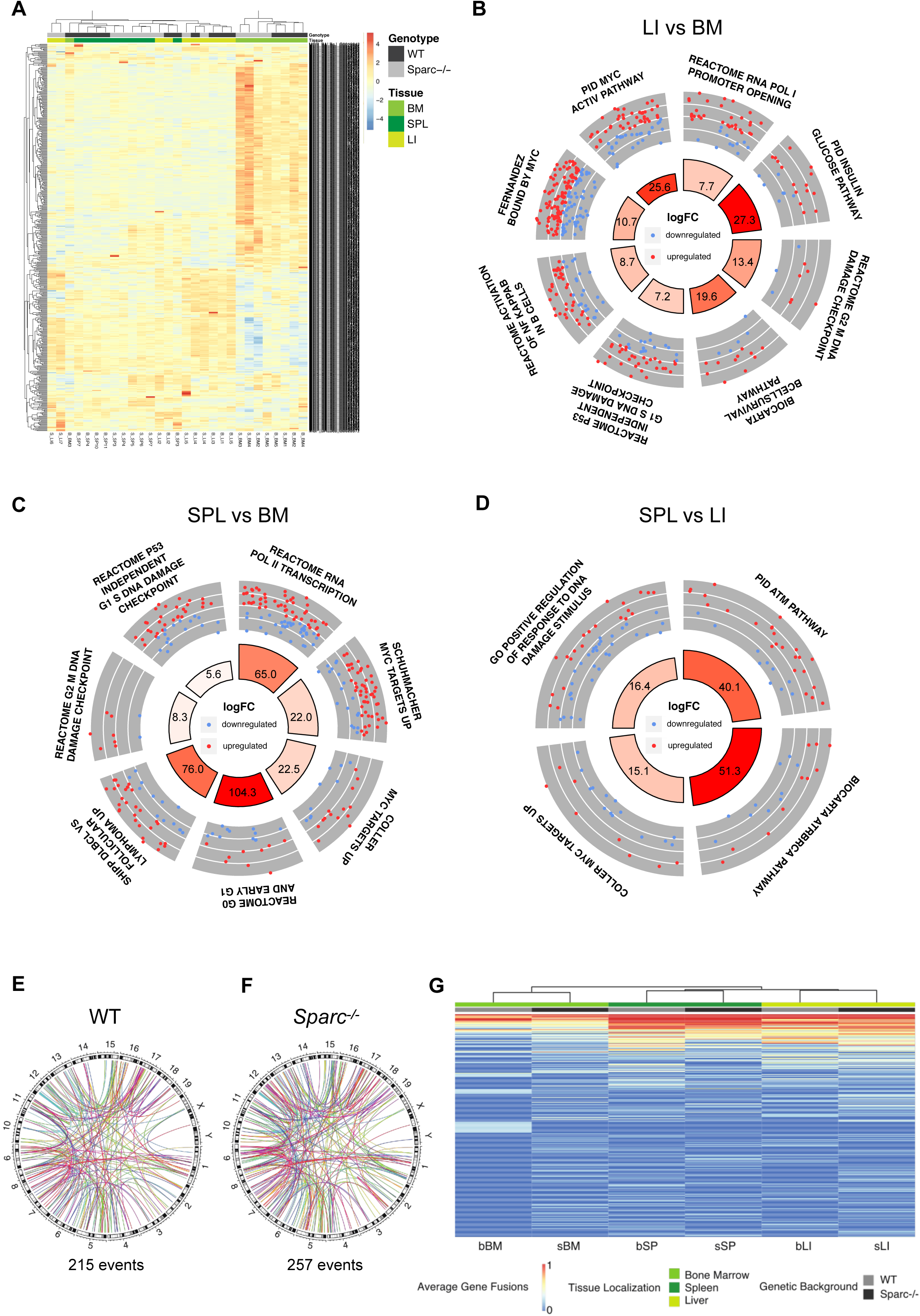
Diversified tissue microenvironment imprints on A20 transcriptional activity. (A) Unsupervised hierarchical clustering of the samples according to the expression of the 500 most variable genes among the samples, showing the tissue imprint on clustering. (B-D) Circular layout visualization of the modulation of selected pathways in LI vs BM (B), SPL vs BM (C) and SPL vs LI (D). The height of the bars of the inner ring represents the significance of the term calculated as -log(p-val). The number within each bar refers to the logFC of the gene set. The outer ring displays scatterplots of the logFC for the genes of the pathway. (E-F) Circular layout visualization of the genomic location of the predicted gene fusions (see Supplemental Methods) in WT (J) Sparc-/- (K) mice. The 19 autosomal and the two sexual mouse chromosomes are shown with the relative chromosome bands. Each colored line represents a fusion event. The number of fusion events is reported under each plot. (G) Unsupervised hierarchical clustering of the gene fusion fraction calculated among all the samples of each genetic background and tissue localization showing a tissue-based imprint on clustering of fusion transcripts (see Supplemental Table S4).

To gain insight into the transcriptional programs modulated among different tissue environments irrespectively of the *Sparc* genotype, we compared the transcriptome profiles of the A20 cells isolated from different tissues by applying Quantitative Set Analysis of Gene Expression (QuSAGE).^22^ The comparison between LI and BM A20 lymphoma cell transcriptional profiles revealed differences in the expression of programs related with metabolism and transcription (Supplemental Table S3, Supplemental Figure S3C), and a significant enrichment in pathways related with B-cell survival and activation of Nf-KB in B cells, as well as in pathways related with DNA damage checkpoints (Figure 3B) in A20 cells purified from LI. Moreover, LI versus BM A20 transcriptomes showed enrichment in transcriptional targets of Myc, including genes identified by ChIP as high-affinity targets of Myc (Figure 3B), pointing to a higher activation of Myc in LI as compared with BM A20 cells. Similarly, the comparison between SPL and BM A20 transcriptomes revealed a significant positive enrichment in pathways involved in transcription, DNA damage checkpoint and Myc transcriptional targets (Figure 3C) in A20 cells purified from the SPL (Supplemental Table S3, Supplemental Figure S3D). Of note, SPL A20 transcriptomes also proved to be enriched in genes reported to be upregulated in DLBCL versus follicular lymphomas^23^ (Figure 3C), which mainly comprised genes involved in glycolytic metabolism, supporting a higher transcriptional activation of programs related with a more aggressive biology. In the comparison between SPL and LI A20 transcriptomes, differences in the transcriptional activation of DNA damage-associated programs were observed, with ATM- and ATR-related pathways being more active in SPL A20 infiltrates (Figure 3D). SPL A20 profiles were also positively enriched in Myc transcriptional targets suggesting that the SPL environment better matched a mutator phenotype (Figure 3D). At contrast, LI A20 transcriptomes showed a positive enrichment in B-cell survival programs, which also characterized the comparison with the BM (Supplemental Table S3, Supplemental Figure S3E).

These results indicate that the BM LI and SPL microenvironment that we found to be characterized by different stromal adaptations to a same DLBCL infiltration, variably promoted or restrained the expression of transcriptional profiles relevant to DLBCL cell biology.

As a control on the effect of transcripts potentially derived from recipient host cells in the analyses, we considered fusion transcript events as a proxy of A20 DLBCL cell specific gene expression. We thus investigated whether a microenvironment-driven clustering of expression data also occurred when only A20 lymphoma-cell specific aberrant gene fusion transcripts were considered. To this aim, fusion events were detected in RNA-seq data (see Supplemental Methods section; Supplemental Table S4), and the distribution and frequency of the fusion events among different sites in the WT and *Sparc*^*-/-*^ genotypes were analyzed (Figure 3E-F). Also in this setting, a tissue microenvironment-driven clustering emerged, sporting fusion events that were more or under-expressed in A20 transcriptomes from one tissue site and/or host *Sparc* genotype, and other fusion events that were expressed at similar levels in all the microenvironments (Figure 3G).

### Myc is modulated in DLBCL cells at mesenchymal interfaces

Given the observed heterogeneity in the A20 lymphoma expression of *Myc* transcriptional programs in microenvironments characterized by different stromal adaptations, we addressed whether *Myc* transcription in A20 cells could be influenced by different mesenchymal elements in *in vitro* co-culture experiments. To this end, A20 cells were cultured in the presence of total Cd45-Cd34-Cd31-Ter119-cKit-Cd29+Sca-1+CD44+ BM-MSCs or An2+ (pericytic) mesenchymal cell fraction (An2+MSCs) from WT and *Sparc*^*-/-*^ mice, and *Myc* expression in A20 cells was assessed at day 2 and day 4 (Figure 4A). A20 cells co-cultured with BM-MSCs showed no significant variation in the expression of *Myc* at day 2 in comparison to basal expression levels (Figure 4B). An induction of *Myc* was observed at day 4 in co-cultures with WT MSCs, and such an induction was not observed in the presence of *Sparc*^*-/-*^ BM-MSCs (Figure 4B). Strikingly, the expression of *Myc* was significantly downmodulated as compared with basal conditions, in co-cultures with both WT and *Sparc*^*-/-*^ An2+MSCs at day 2 and 4 (Figure 4B). This finding hinted to a potential influence of stromal neighbors over the expression of *Myc* in such high-grade lymphomatous cells.

**Figure 4.**
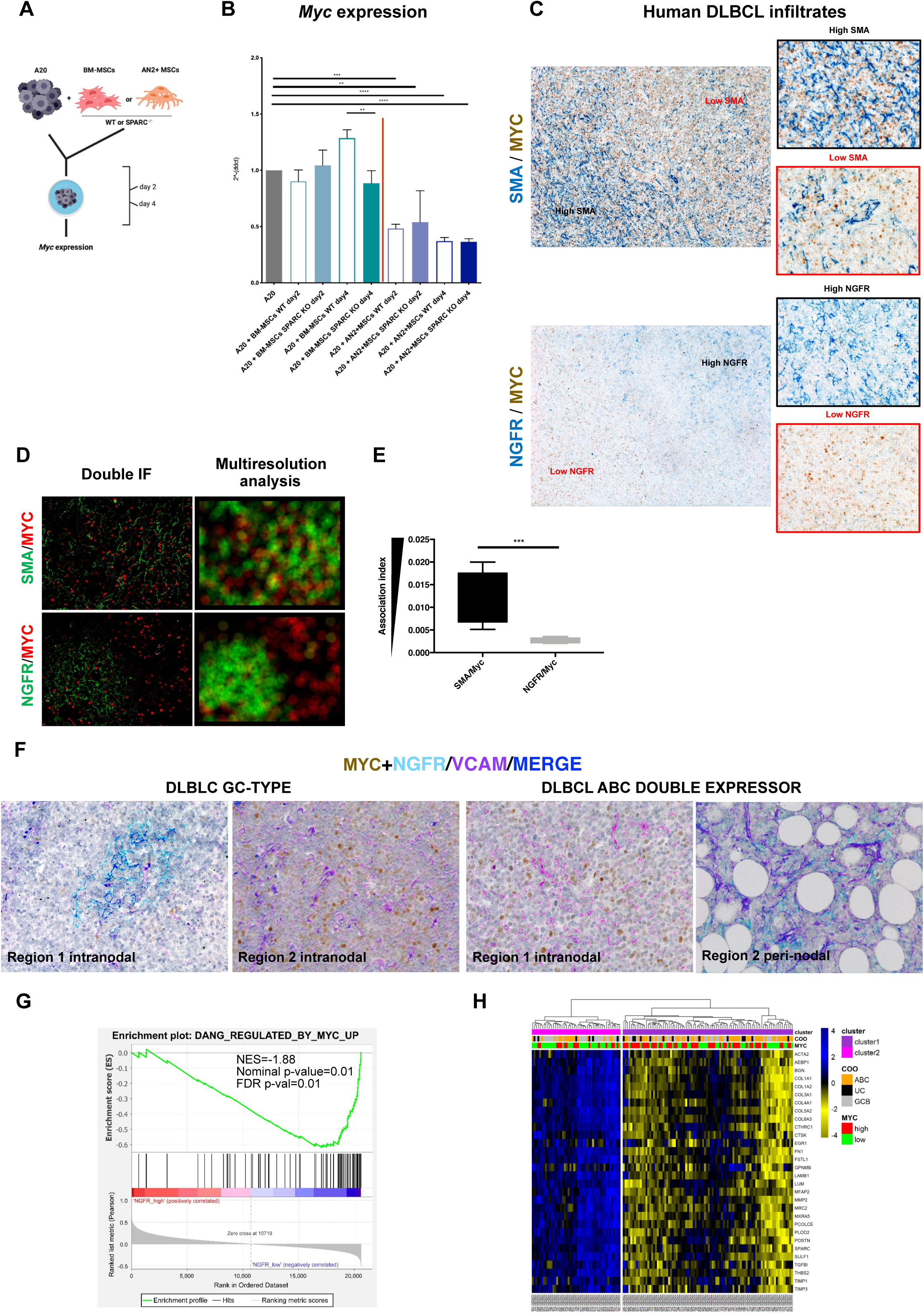
Different mesenchymal contextures correlate with Myc modulation and spatial heterogeneity. (A) Graphical scheme of the in vitro co-culturing experiment with A20 cells and either WT or *Sparc*^*-/-*^ total BM-derived MSCs or peri-vascular An2+MSCs (see Methods section for MSC purification and phenotyping). (B) Myc gene expression by Real-time PCR on total RNA isolated from A20 cells at basal conditions and following 2 or 4 days co-culture with WT or *Sparc*^*-/-*^ BM-MSCs or An2+MSCs, showing the significant downmodulation of Myc in the presence of the latter mesenchymal cell phenotype, independently of MSC *Sparc* status. (C) Double-marker IHC for SMA (blue signal, top panels) and MYC (brown signal) or NGFR (blue signal bottom panel) and MYC (brown signal) on human DLBCL lymph node infiltrates (Supplemental Table S1) indicating that no MYC expression gradient is evident between SMA-high and SMA-low areas, while an inverse distribution of MYC and NGFR density emerges. Original magnifications x50 and x400 (insets). (D) Representative input (left panels) and output (right panels) images of Multiresolution Analysis (see Supplemental Methods Section) applied to double-marker IF stainings for SMA (green signal, upper panels) and MYC (red signal) or NGFR (green signal, lower panels) and MYC (red signal) on human DLBCL samples (Supplemental Table S1), showing the spatial segregation of NGFR- and MYC-stained areas (nearly complete absence of yellow signal in the output panel) and the intermingling SMA and MYC stainings (appearance of yellow signal in the output panel). Original magnification x200. (E) Graphicaon output of the Association index calculated according to the Multiresolution Analysis applied to SMA/MYC and NGFR/MYC immunostainings on human DLBCL samples (Supplemental Table S1) and expressed as the inverse of the average distance between the signals corresponding to the protein markers. The output shows a significantly lower degree of association between NGFR and MYC. (F) Representative multiplexed chromogenic and fluorescence immunostainings for MYC (brown signal), NGFR (cyan signal), and VCAM-1 (violet signal) on human DLBCL samples (Supplemental Table S1) showing the different expression of MYC among areas with higher or lower NGFR expression. VCAM-1 highlights the presence of a stromal cell network also in areas devoid of NGFR. Original magnification x200. (G) GSEA output showing the positive enrichment of MYC-upregulated target genes, in human DLBCL cases characterized by lower (above median) NGFR expression, using GSE98588 dataset (NES, normalized enrichment score; FDR, false discovery rate). (H) Heatmap related to the expression of the 30 genes characteristic of stromal compartment in DLBCL from GSE98588 dataset. Signal intensities ranked from highest (blue) to lowest (yellow) are indicated as row z-score. On the top, information including clusters (derived from hierarchical clustering analysis), cell of origin (COO) and MYC expression are depicted. The analysis shows that stromal gene-based clustering of the 137 cases into two groups determines differential enrichment of MYC-high and -low cases.

Prompted by the observed *Myc* modulation in *in vitro* co-culture experiments, we investigated whether the intra-tumor mesenchymal heterogeneity that we highlighted in the small cohort of human nodal DLBCLs could relate with *in situ* MYC expression. To this aim, we applied an *ad-hoc*-developed software tool for multiresolution spatial analysis (see Supplemental Methods section) to immunostained sections for the myofibroblastic/reticular cell marker SMA and for the MSC/pericytic marker NGFR in combination with MYC, and evaluated the degree of spatial association between the two mesenchymal phenotypes and MYC expression in lymphomatous infiltrates. Interestingly, while SMA+ stroma distribution did not show a significant degree of association with MYC, NGFR+ mesenchymal foci were negatively associated with MYC expression (Figure 4C-E), suggesting that a topographic compartmentalization relative to mesenchymal cell distribution and/or phenotype acquisition could occur. Indeed, in MYC-expressing DLBCLs (>40% of MYC+ cells), areas characterized by an NGFR-rich meshwork showed a significantly lower MYC expression as compared with areas lacking NGFR but still showing a dense stromal meshwork as highlighted by VCAM-1 (Figure 4F). Remarkably, in cases in which the nodal DLBCL proliferation extended to the peri-nodal adipose tissue highly rich in NGFR+ perivascular mesenchymal elements, a neat downmodulation of MYC expression was observed, which provided an insight into the dynamical switch of MYC expression under specific stromal microenvironment conditions (Figure 4F). Consistently with the observed negative association between NGFR+ stroma and MYC expression, GSEA for MYC transcriptional targets on 137 DLBCL cases^4^ categorized according to *NGFR* expression levels into *NGFR*-high and -low showed a significant enrichment of upregulated MYC targets in *NGFR*-low cases (NES=-1.88, FDR=0.01, Figure 4G). Moreover, zooming out on whole transcriptome, we performed hierarchical clustering analysis of 30 stromal genes previously described as specific of the mesenchymal compartment in DLBCL^7^ exploiting the same 137 cases^4^ and observed a significantly different distribution of cases stratified by *MYC* median expression between the two clusters characterized by high or low levels of the 30-gene mesenchymal signature (Fisher’s exact test, p=0.01, Figure 4H), which pointed to clear-cut differences in the mesenchymal milieu of cases with divergent *MYC* expression levels.

Altogether these data point to topographic influences over relevant DLBCL phenotypes such as MYC expression, related with mesenchymal determinants, the mechanisms of which deserve to be thoroughly dissected.

### Stromal *Sparc* proficiency tunes the immune pressure on A20 lymphomas

Following the analysis of transcriptional and phenotypic heterogeneity related with tissue microenvironment-intrinsic determinants, we finally investigated whether Sparc proficiency could also play a role in modulating A20 lymphoma dynamics. The transcriptional differences between A20 cells from WT and *Sparc*^*-/-*^ hosts were mostly contributed by genes differentially expressed between the WT and *Sparc*^*-/-*^ BM samples (Figure 5A), consistently with the prominent expression of Sparc in the BM stroma.^24^ We performed QuSAGE analysis by comparing A20 transcriptomes from WT and *Sparc*^*-/-*^ infiltrates including all tissue sites (Supplemental Table S3, Supplemental Figure S4A). In the *Sparc*^*-/-*^ host, a significantly higher activation of the *Ras* and *Rhoa* GTPase pathways was detected in association with increased activity of Foxo and Glycogenolysis pathways (Figure 5B), suggesting a different type of metabolic adaptation to environment sensing. Moreover, A20 transcriptomes in the Sparc-deficient environment harbored a positive enrichment in genes involved in Complement activation (Figure 5B), which is in line with our previous demonstration of defective stromal Sparc enhancing the expression of cell-intrinsic innate immune/inflammatory programs also involved in tumor-stroma interactions.-^25-26^ Consistently with an environment more prone to immune activation, A20 infiltrates in *Sparc*^*-/-*^ hosts showed a higher density of infiltrating T cells assessed by CD3 and CD8 quantitative IHC, as compared with the WT counterpart (Figure 5C-D). The difference was particularly prominent in the BM microenvironment, suggesting a more profound alteration of the BM immune contexture in the absence of Sparc. To evaluate whether the differences observed in the T cell infiltration of *Sparc*^*-/-*^ and WT BM environment could result in a different immune pressure on lymphomatous cells, A20 (GFP+) cells were injected intravenously into WT and *Sparc*^*-/-*^ hosts and then recovered from the BM for serial i.v. transplants into either WT or *Sparc*^*-/-*^ syngeneic hosts according to the donor genotype (Figure 5E). After 4 passages, A20 cells conditioned by either WT or *Sparc*^*-/-*^ microenvironment were compared for growth into both WT and *Sparc*^*-/-*^ *recipients*, through FACS analysis of A20 cells in the BM and SPL (Figure 5E). Moreover, the expression of MHC class-I and class-II on A20 cell surface and the frequency and activation of CD8+ infiltrating T cells, were analyzed. We observed an increased take of A20 cells conditioned within the Sparc-deficient BM environment (A20-BM*Sparc*^*-/-*^) regardless their final transplant into WT or *Sparc*^*-/-*^ recipients (Figure 5F). This was associated with a significant down-modulation of MHC-I, but not of MHC-II, on the cell surface of A20-BM*Sparc*^*-/-*^ compared to A20-BMWT (Figure 5G-H). Although the frequency of total CD8+ T cells only barely differed among BM samples infiltrated by A20-BM*Sparc*^*-/-*^ and A20-BMWT (Figure 5I), the frequency of activated proliferating CD8+PD-1+Tim-3-Ki-67+ T cells was lower in the presence of A20-BM*Sparc*^*-/-*^ cells (Figure 5J), indicating a lower degree of effector T cell activation in samples with lower MHC-I levels. Notably, in the same samples, a lower frequency of CD4+CD25+Foxp3+ regulatory T cells was detected (Figure 5K), further supporting the selection of a less immunogenic profile in the *Sparc*^*-/-*^ BM microenvironment. The extremely low fractions of GFP+ A20 cells detected in the SPL of the same hosts (Supplemental Figure S4B) indicated that BM microenvironment priming of A20 cells mainly fostered BM infiltration, in line with the centrality of tissue environment-specific determinants in the stromal/immune adaptation to lymphoma growth.

**Figure 5.**
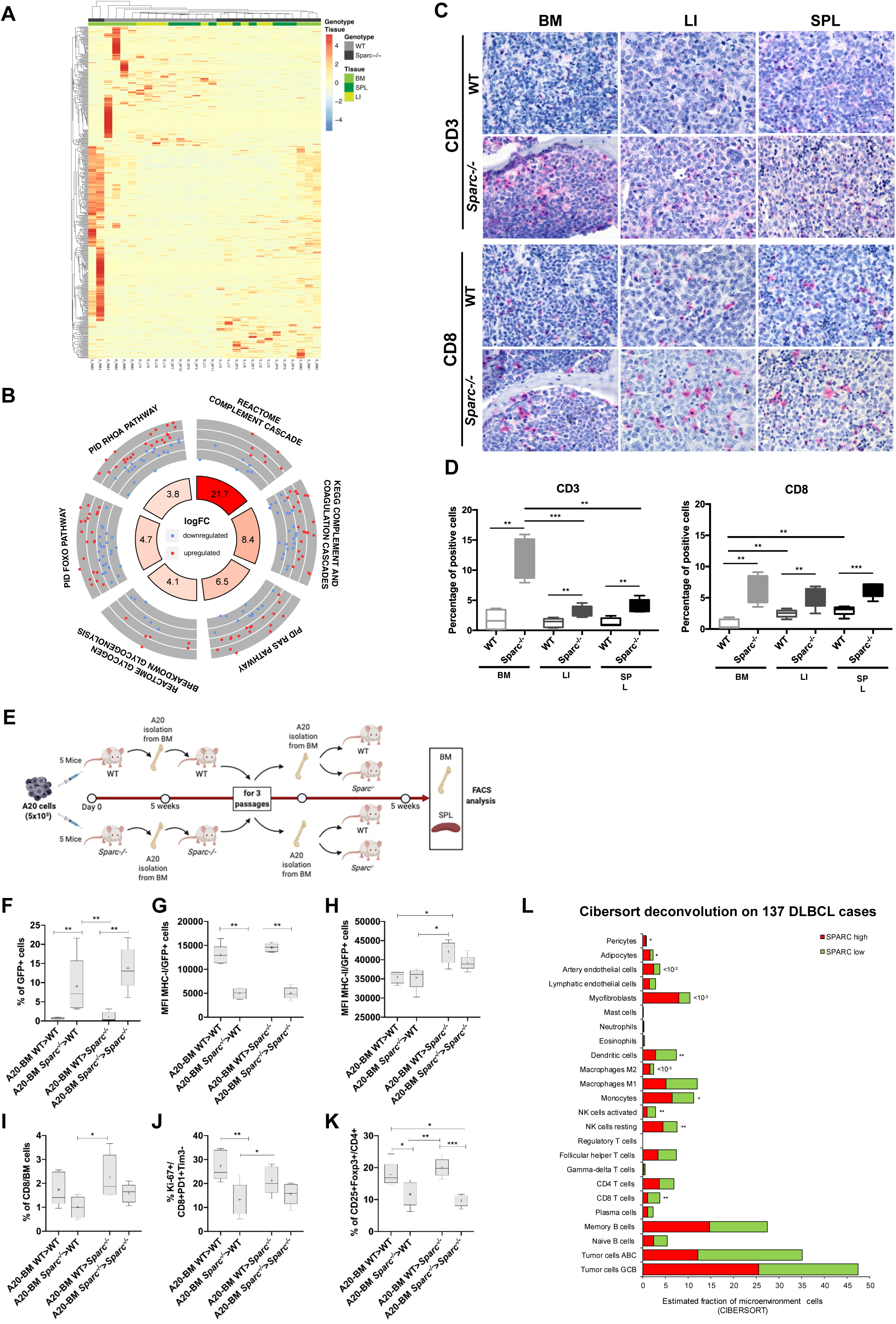
Stromal Sparc status conditions immune pressure and escape. (A) Heat map showing the unsupervised hierarchical clustering of the samples according to the top 500 deregulated genes between A20 lymphoma cells injected in WT and *Sparc*^*-/-*^ mice. Genes were ranked according to absolute value of their log2FC between cells isolated from WT and *Sparc*^*-/-*^ tissues. The heatmap highlights the prominent contribution of BM samples to WT vs *Sparc*^*-/-*^ differential gene expression. (B) Circular layout visualization of the differences in the activities of genes belonging to gene sets involved in Ras, Rhoa, Foxo signaling pathways, glycogenolysis, and Complement cascade, between A20 cells isolated from WT and *Sparc*^*-/-*^ mice as assessed by QuSAGE. The height of the bars of the inner ring represents the significance of the term calculated as -log(p-val). The number within each bar refers to the logFC of the gene set. The outer ring displays scatterplots of the logFC for the genes of the pathway. (C) IHC for CD3 (upper panels) and CD8 (lower panels) in BM LI and SPL A20 DLBCL infiltrates of WT and *Sparc*^*-/-*^ mice, highlighting significant differences in the density of infiltrating T cells in the two stromal settings. Original magnification x400. (D) quantitative analysis of CD3 and CD8 IHC in BM, LI, and SPL A20 DLBCL infiltrates in WT and *Sparc*^*-/-*^ mice. (E) Graphical scheme of the in vivo experiment of A20 injection, BM purification and re-transplantation into WT and *Sparc*^-/-^ hosts, aimed at investigating the effects of immune priming by WT and *Sparc*^*-/-*^ BM microenvironment on A20 lymphomas. (F-K) Flow cytometry analyses of the infiltrating A20 (GFP+) cell fraction (F); MHC class-I (G) and class–II (H) mean fluorescence intensity (MFI) on A20 (GFP+) cells; total (I) and activated (J) proliferating CD8+ T cell effectors and regulatory T cells (K), in the BM of WT and *Sparc*^*-/-*^ final recipients of A20 cells conditioned by three passages into WT and *Sparc*^*-/-*^ BM. (L) CIBERSORT deconvolution analysis output of the relative frequencies of immune and stromal cell populations according to the levels of SPARC expression in the GSE98588 dataset of human DLBCLs, showing that SPARC-low cases are significantly enriched in CD8, activated NK, and Dendritic cell populations, as compared with SPARC-high cases.

Based on the *in vivo* results pointing to a stronger immune pressure exerted on A20 lymphomatous infiltrates when in a Sparc-defective stromal setting, we wanted to investigate whether human DLBCL cases characterized by different *SPARC* expression levels could display signs of a different T cell infiltration. We applied the CIBERSORT deconvolution algorithm (see Methods section) to the transcriptomes of the 137 DLBCL dataset adopted for *MYC* GSEA analysis,^4^ using a customized gene signature matrix generated as previously reported^7^ and detailed in the Methods section. The CIBERSORT output revealed that CD8 T cells, activated NK cells and Dendritic cells frequencies were significantly higher in low-*SPARC*, as compared with high-*SPARC* DLBCL cases (Figure 5L, Supplemental Table S5). This evidence further underlines the relationship between the control of immune activation and *SPARC* expression in the DLBCL microenvironment, suggesting that the reported unfavorable prognostic predicted by low *SPARC* may be to some extent expression of a more efficient immune editing.

## DISCUSSION

The current challenge in DLBCL is represented by the identification of mechanisms for standard chemo-immuno treatment failure in *bona fide* prognostically favorable GC-related DLBCLs, and by the identification of successful strategies for cases in which strong drivers of DLBCL aggressiveness, such as MYC and BCL2 overexpression negatively impact on the disease course. The recent focus on the integration among mutational landscape and gene expression profiles identified DLBCL subsets independent from the cell of origin, with distinct mutational trajectories and druggable pathways, including clusters identified by recurrent EZH2 and other chromatin modifier gene mutations or by genetic lesions in JAK/STAT and NF-kB pathways.^4-5^ Yet, these achievements did not shed light on the possible transcriptional and/or phenotypic heterogeneity that is coded at the topographic level. We described here, in a syngenic model of DLBCL, that the expression of biologically relevant features, such as Myc and its target genes, and the modulation of DNA damage-associated programs are consistently influenced by the tissue microenvironment. This is in line with the recently reported evidence of diversified subclonal complexity within lymphoid and non-lymphoid tissues in spontaneous Myc-driven lymphomas, as a result of immune editing.^27^ The differences in the stromatogenesis observed in the BM, LI, and SPL A20 DLBCL tumors point to a still neglected contribution of resident mesenchymal populations to the intra-tumor diversification of these high-grade malignancies. In solid cancers, specifically in pancreatic adenocarcinoma, cancer-associated fibroblasts have been recently demonstrated to modulate the acquisition of proliferative and/or epithelial-to-mesenchymal transition phenotypes in malignant cells, shaping intra-tumor heterogeneity within the glandular component.^28^

Along with the tissue-site-related modulations of A20 transcriptomic and phenotypic features, we herein also identified heterogeneous mesenchymal foci within the same human DLBCL nodal lesions, which related differently to MYC expression in lymphomatous cells. Within secondary lymphoid organs, resident mesenchymal cells represent a highly diversified constellation of non-hematopoietic elements that finely tune the functional differentiation and activity of the immune cells taking part into the ongoing immune responses.^29^ Among these populations, FDCs and perivascular stromal cells display high phenotypical homology with bone marrow-derived mesenchymal stromal cells expressing NGFR and CD146 MSC markers.^10,30-31^ We found that DLBCL infiltrates characterized by the presence of NGFR+ mesenchymal foci displayed MYC downregulated expression within the same areas, as compared with areas characterized by a predominance of other mesenchymal phenotypes such as SMA+ and/or VCAM-1+ reticular elements. This finding is apparently conflicting with the physiological expression of MYC by a fraction of B cells within the light zone of the GC, which features a dense NGFR+ FDC meshwork, and suggests that the normal dynamics of B-cell/stroma competition may be profoundly altered along malignant transformation, a concept that is recently emerging in human cancer.^32^ The finding that the MYC heterogeneous expression may associate with diversified stromal interfaces claims for a consideration regarding the actual accuracy of the determination of MYC expression in the diagnostic algorithm of DLBCL ^33^ and allows speculating on the possibility of different paces in the disease progression and/or treatment response influenced by topographic stroma determinants.

In our experiments we also adopted *Sparc*^*-/-*^ hosts in the attempt to gain insight into the biological bases of the reported stromal Sparc prognostic influence in DLBCL. Indeed, within a signature of 30 stromal genes correlated with prognosis in haematological malignancies,^9^ *SPARC* showed the strongest association with a favorable prognosis in the DLBCL setting (Supplemental Table S6). The positive prognostic influence cast by SPARC has been correlated with its myeloid cell-intrinsic expression^34^ and related with the ABC setting, in which oncogenic activation of inflammatory pathways has been described.^35^ Consistently with SPARC influencing the tumor inflammatory microenvironment through a myeloid cell-related function, we have previously demonstrated that the abundance of Sparc in the stromal milieu of cancer dictates the pro-inflammatory or suppressive phenotype of infiltrating myeloid cells.^26^ In the present A20 model, we observed that lymphomatous cells residing in Sparc-defective environment showed transcriptional activation of pathways related with Complement cascade triggering and activation. This finding is consistent with the preferential induction of Complement recognition factor C1q and C5a effector expression in the BM microenvironment of mice with genetically-driven myeloproliferation in the absence of stromal Sparc.^36^ Along with immune effector and co-stimulation, Complement factors may exert non-canonical functions in the tumor microenvironment, including tumor-promoting roles through enhancement of tumor-stroma interactions, angiogenesis and chemotaxis of tumor-associated myeloid cells.^37-40^ Notably, activation of Complement recognition factors in the tumor setting hardly results in complete activation of the Complement cascade with unleashing of its cytotoxic function because of the ubiquitous expression of Complement regulatory proteins (i.e. CD46, CD55, CD59) on the cell surface and on the expression of intermediate Complement factors as decoy of membrane attack complex formation.^41-42^ Among other pathways induced in A20 transcriptomes in the *Sparc*^*-/-*^ setting, were those associated with *Ras, Rhoa* and *Foxo* activity, which have been implicated in the pathogenesis of ABC-DLBCL.^43-44^ Modulation of Ras, Rhoa, and Foxo1 signaling is strongly implicated in BCR activity,^45-46^ which has been reported to crucially influence the fitness of Myc-driven high grade B-cell clones.^47^ The observed difference in the trancriptional activity of these key pathways could indicate different assets of environment sensing and response by A20 cells embedded in WT and *Sparc*^*-/-*^ stroma. Further corroborating the hypothesis of a tumor/stroma interface altered by Sparc deficiency, A20 infiltrates in the *Sparc*^*-/-*^ setting showed a denser degree of T-cell infiltration. These results found correlative evidence in human DLBCL transcriptional deconvolution analyses by CIBERSORT, which indicated a significant enrichment in CD8 and NK activated effectors in the setting of low *SPARC* expression. Moreover, upon multiple passages in *Sparc*^*-/-*^ or WT BM, A20 cells were differently enriched in MHC-I-negative populations in a way that was dependent on the genotype of the priming environment, indicating that a stronger selective pressure towards immune escape was exerted by the *Sparc*^*-/-*^ microenvironment. This evidence adds a piece of information to the recent characterization of the genetic landscape of MHC deficiency in DLBCL,^48^ suggesting that a level of regulation exerted by the stromal microenvironment may intervene determining topographic biases in MHC-negativity selection.

In conclusion, our study provides experimental evidence that stromal microenvironment generates topological determinants of intra-tumor heterogeneity in DLBCL that involve key transcriptional pathways potentially conferring spatial segregation to prognostically relevant features, such as Myc expression and/or DDR programs. We envisage the upcoming emergence of new classification/prognostication efforts based on the representation of spatially-resolved mesenchymal-immune ecosystems.

## ACKNOWLEDGEMENTS

This Study has been supported by the Italian Foundation for Cancer Research (AIRC) (grants 15999 and 22145 to C. Tripodo) and by the University of Palermo. The Authors acknowledge Dr Stefano Casola (IFOM, the FIRC Institute for Molecular Oncology) for expert advice on experimental design and results interpretation, and Dr Simona De Summa (Giovanni Paolo II Cancer Institute, Bari) for helpful discussion on bioinformatics analyses. The Authors are also grateful to Dr Nadia Castioni for technical support.

## AUTHORSHIP

### Contribution

S. Sangaletti, F. Iannelli, F. Zanardi: Conceptualization, Data curation, Formal analysis, Investigation, Visualization, Methodology, Writing. V. Cancila, P. Portararo, L. Botti, D. Vacca, A. Di Napoli, C. Chiodoni, F. Pisati, A. Gulino, M. Ponzoni: Resources, Data curation, Formal analysis, Investigation, Visualization, Validation, Writing C. Valenti, M. Vegliante: Data curation, Software, Formal analysis, Investigation, Methodology, Writing. M.P. Colombo, C. Tripodo: Conceptualization, Data curation, Formal analysis, Supervision, Funding acquisition, Validation, Investigation, Visualization, Methodology, Writing.

