## Supplemental Methods and Figures for "Intra-tumor heterogeneity of Diffuse Large B-cell Lymphoma involves the induction of diversified stroma-tumor interfaces"

**Generation of GFP+ A20 cells through GFP-expressing lentiviral vector**

Lentiviral vector pRRLCMVGFPsin-18 (a kind gift from Dr. G. Ferrari from San Raffaele Hospital, Milan, Italy) has been used to transduce A20 cells. A third-generation packaging system was used to produce viral particles. Lentiviral stocks were produced in 293T cells by Ca3PO4 co-trasfection of the plasmids as previously described (1). A20 cells were infected with lentiviral particles added at a multiplicity of infection (MOI) of 20. After 24 hours, cells were washed, and expanded. Three days later cells were checked for efficiency of transduction. To obtain a pure population of GFP-expressing cells, we sorted GFP+ A20 cells using a FACSAria sorter (BD).

**Immunophenotypical analyses**

For immunophenotypical analyses, human and mouse tissue sections were deparaffinized, rehydrated and unmasked using Novocastra Epitope Retrieval Solutions pH 6 and pH 9 in PT Link Dako at 98°C for 30 minutes. Subsequently, the sections were brought to room temperature and washed in PBS. After neutralization of the endogenous peroxidase with 3% $H_2O_2$ and Fc blocking by a specific protein block (Novocastra), the samples were incubated with antibodies. The following primary antibodies were adopted for IHC and IF: Ki67 (Abcam, 1:1000 pH6), c-Myc (clone Y69 Abcam, 1:500 pH6), p-γH2ax (Abcam, 1:1000 pH6), p-Chk1 (Abcam, 1:500 pH9), CD3 (Abcam, 1:100 pH9), CD8 (clone D4W2Z Cell Signaling, 1:400 pH9), αSMA (clone 1A4(asm-1 Biocare, 1:500 pH9), NGFR (clone 7F10 Leica Microsystems, 1:50 pH6), VCAM1 (clone EPR5047 Abcam, 1:500 pH9), CD146 (clone N1238 Abcam, 1:25 pH9), CD146 (clone EPR3208 Abcam, 1:25 pH9), PDGFR-β (clone Y92 Abcam, 1:250 pH6), Nestin (clone rat-40 1 Millipore, 1:100 pH9). Anti-mouse and anti-rabbit (Alexa Fluor 488 and 568 Conjugate) secondary antibodies were used for fluorescence detection.
IHC staining was revealed using Novolink Polymer Detection Systems (Novocastra) or IgG (H&L) specific secondary antibodies (Life Technologies, 1:500) or MACH 2 Double Stain 1 kit and DAB (3,3'-Diaminobenzidine, Novocastra) or Vulcan Fast Red or Ferangi Blue as substrate-chromogens.

**Multi-resolution analysis**

For multiresolution analysis of double-marker immunostained tissue sections, an efficient convolution of the redundant "à trous" algorithm with bilinear spline interpolation kernel has been realized; this has guaranteed an isotropic filter that therefore does not prefer particular directions (i.e. shapes). For the 20x and 40x magnifications, planes 5 and 6 of the dyadic analysis were chosen respectively, since they experimentally showed the best automatic segmentation with simple binarization on threshold equal to the mean value of the wavelet coefficients, increased by 1.5 times their standard deviation. This technique is described in literature for a rapid and correct segmentation, while attenuating eventual noise. Among the measures we considered (e.g. Pearson statistical index, Structural Similarity index, Peak Signal-to-Noise Ratio), the Euclidean distance between the centroids of the two classes (SMA/MYC and NGFR/MYC) was effective in underlining the three conditions of co-location, non-localization and substantially random distribution.

**Flow cytometry**

For flow cytometry analyses, the following antibodies were used: I H-2K$^d$ (PE, clone SF1-1.1.1, eBioscience), I-A/I-E (APCeFluor780, clone M5/114.15.2, eBioscience), CD134 (OX-40) (PE, clone OX-86, Biolegend), CD278 (ICOS) (BV510, clone C398.4A, Biolegend), CD8 (BV605, clone 53-6.7, BD), CD25 (BV650, clone PC61, BD), CD279 (PD-1) (BV786, clone J43, BD), Foxp3 (PerCpCy5.5, clone FJK-16.5, eBioscience), Ki-67 (APC, clone SolA15, eBioscience), CD4 (APC-Cy7, clone RM4-5, Biolegend),

CD366 (TIM3) (BV421, clone 5D12/TIM-3, BD), CD45 (FITC, clone 30-F11, Tonbo Biosciences), CD31 (FITC, clone 390, eBioscience), TER119 (FITC, clone Ter119, Invitrogen), CD34 (FITC, clone RAM34, BD), cKIT (FITC, clone 2B8, eBioscience), SCA-1 (APC, clone D7, BD), CD44 (APC-Cy7, clone IM7, BD).

**RNA sequencing and data analysis**

The sample B-BM1 was removed from the analyses due to the low number of mapped reads (~6.7 million reads) and following the evaluation of the Euclidean distance among the samples. Differentially expressed genes were called using DESeq2 R package (2) starting from the count tables generated by HTSeq (3).

Quantitative gene set analysis was performed with the QuSAGE R package (4) using the log2 normalized expression values and either the c2 curated gene sets of the Broad Molecular Signatures Database (MSigDB; http://software.broadinstitute.org/gsea/msigdb/collections.jsp) or a custom list of gene sets manually selected for being implicated in DNA damage, MYC targets or B-cell biology (Supplemental Table S7).

Heatmaps representing the unsupervised hierarchical clustering of the samples according to the log2 normalized expression values of genes belonging to the gene set of interest were generated with the function pheatmap of the pheatmap (v 1.012) R package.

Circular layout visualization of the logFC of the activity of selected pathways were generated with a custom R script that made use of the GOplot (5) R library.

Gene fusion events were detected in RNA-seq data using the tool FusionCatcher (https://github.com/ndaniel/fusioncatcher), and *ad-hoc* R scripts were used to process the data in order to remove possible false positives according to FusionCatcher "annotation" field that takes into account several factors like known false positives, fusion genes seen in healthy sample or paralogy of genes involved in the fusion event (see FusionCatcher manual) (Supplemental Table S4). Circular layout visualization of the genomic location of the predicted gene fusions in WT and *Sparc-/-* samples were generated with a custom R script based on the circlize R package.

#### SUPPLEMENTAL FIGURE LEGENDS

**Supplementary Figure S1 Flow cytometry-based forting of A20 cells from tissues.**
(A) Representative flow cytometry sorting of A20 cells from BM, Spleen (highly and lowly infiltrated) and Liver of mice injected with A20 cells accrding to the experiment's graphical scheme shown in Figure 1A. Within the SSC-high CD45+ gate, A20 cells were characterized for high B220 and CD19 expression.

**Supplemental Figure S2 Intralesional heterogeneity of the mesenchymal components in DLBCL infiltrates.**
(A) IHC for SMA, NGFR, CD146, and PDGFRb mesenchymal markers on three different regions (coloured insets) from two representative human DLBCL (see Supplemental Table 1) lymph node biopsies (panoramic insets) highlighting the disconnected staining pattern of the four markers in the three regions, which indicates the existence of different stromal adaptations to lymphoma infiltration within the same lesion. Regions were selected from whole slide digital scans. Original magnifications, x2 (panoramic insets) and x200 (coloured insets). (B) double marker IF for NGFR (green signal) and either SMA, PDGFRb, or VCAM-1 (all in red), on lymphoid follicles from a human lymh node with reactive follicular hyperplasia (upper panels) or infiltrates from a human DLBCL lymph node biopsy (lower panels). Mesenchymal markers that identify the FDC meshwork (NGFR and VCAM-1) or the inter-follicular fibroblastic reticular and sinusoidal network (SMA and PDGFRb) in the reactive condition (upper panels) are aberrantly focally associated or dissociated in DLBCL infiltrates (lower panels, arrows). Original magnification, x200.

**Supplemental Figure S3 Principal component analysis and clustering of the A20 transcriptomes from BM, LI, and SPL, and quantitative set analysis for gene expression.**
(A) Principal component analysis of the samples. (B) Unsupervised hierarchical clustering of the samples according to their Euclidean distance calculated on the rlog-transformed data. (C-E) Bar plots displaying the 100 most modulated pathways between LI and BM (C), SP and BM (D) and SP and LI (E), as assessed by QuSAGE. Each bar shows the mean and the confidence interval of the gene set.

**Supplementary Figure S4 Quantitative set analysis for gene expression in WT versus Sparc-/- comparison and lymphoma cell frequency in the SPL of WT and *Sparc-/-* mice upon transplantation of A20-GFP+ cells.**
(A) Bar plots displaying the 100 most modulated pathways between WT and *Sparc-/-* samples (including BM, LI and SPL tissue sites), as assessed by QuSAGE. Each bar shows the mean and the confidence interval of the gene set. (B) The frequency of A20 cells infiltrating the SPL of WT and *Sparc-/-* mice has been evaluated after 5 weeks of the injection of $5x10^5$ A20-GFP+ cells from the BM, passaged three times in WT or *Sparc-/-* hosts as detailed in the experiment's graphical scheme in Figure 5E. The SPL frequency of A20-GFP+ cells is extremely low and does not significantly vary according to the *Sparc* genotype of the priming or recipient hosts.

### SUPPLEMENTAL FIGURES

#### Supplemental Figure S1

**A**

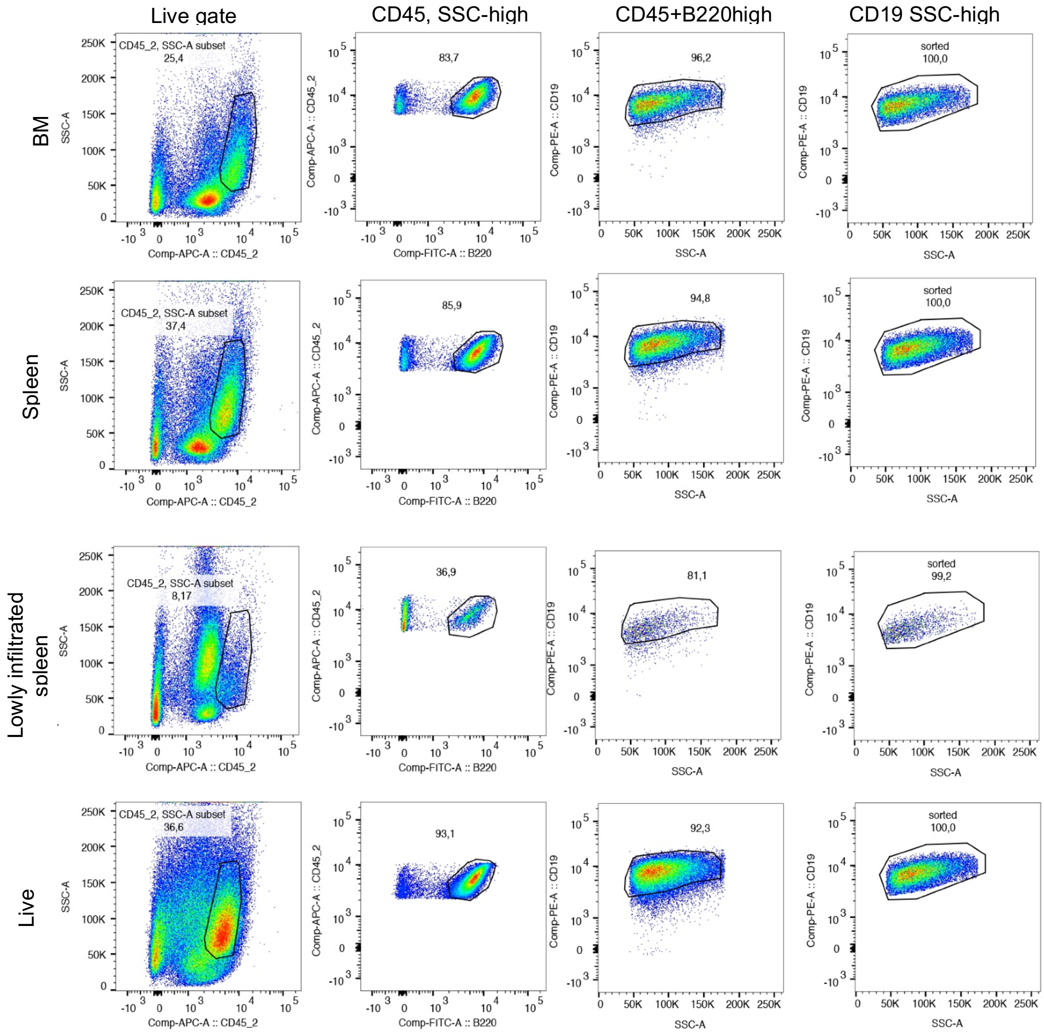

Supplemental Figure S2

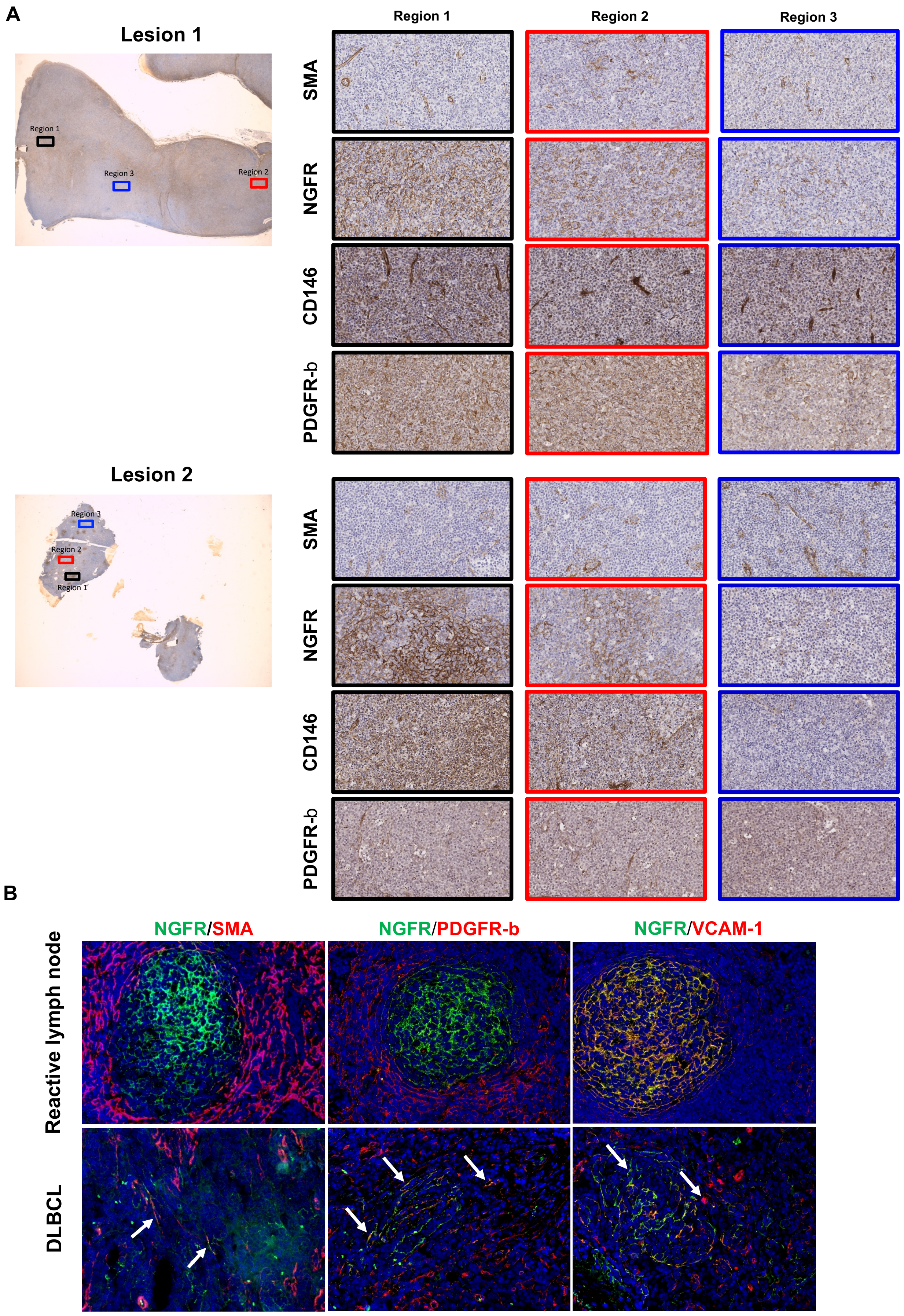

#### Supplemental Figure S3

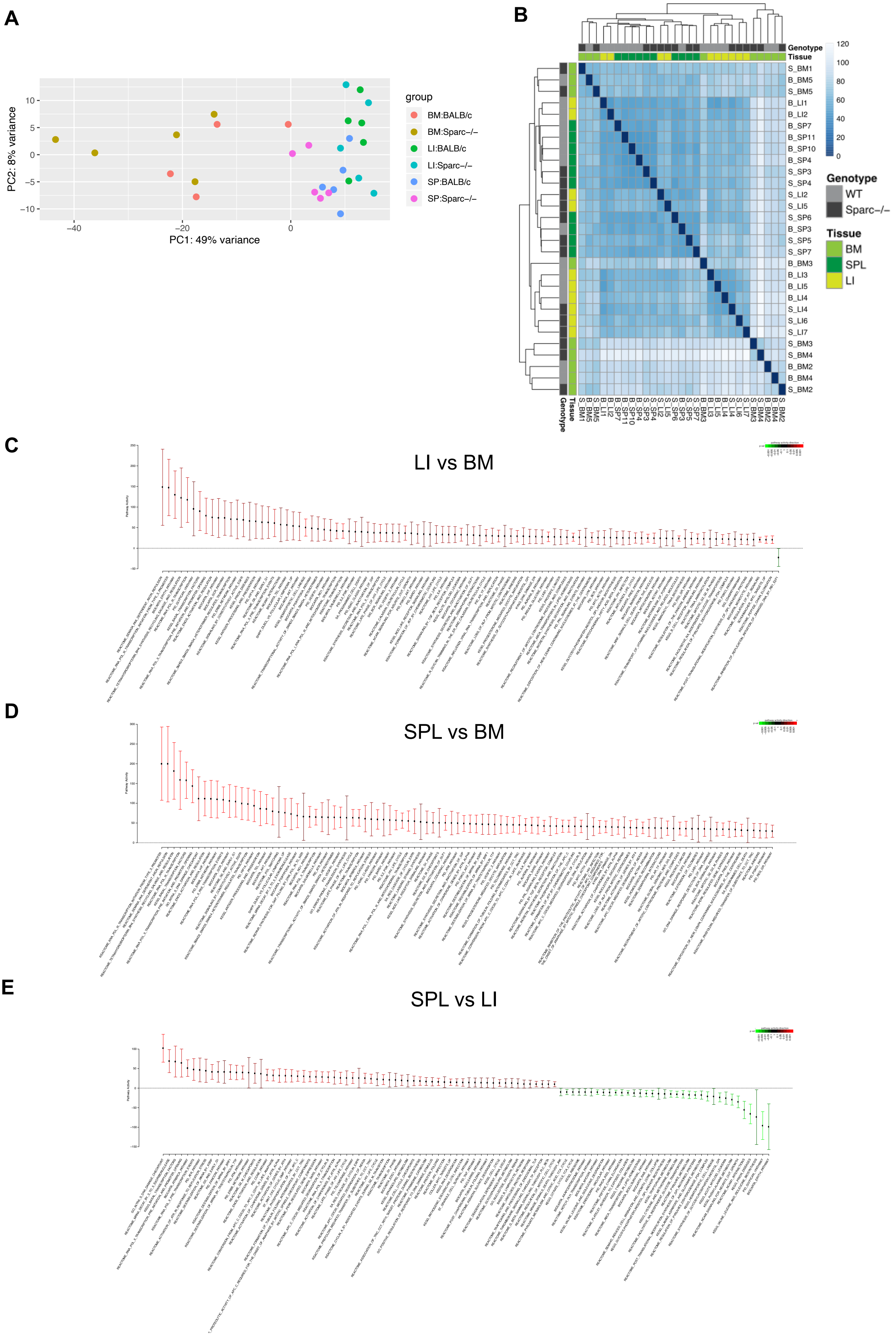

#### Supplemental Figure S4

**A**

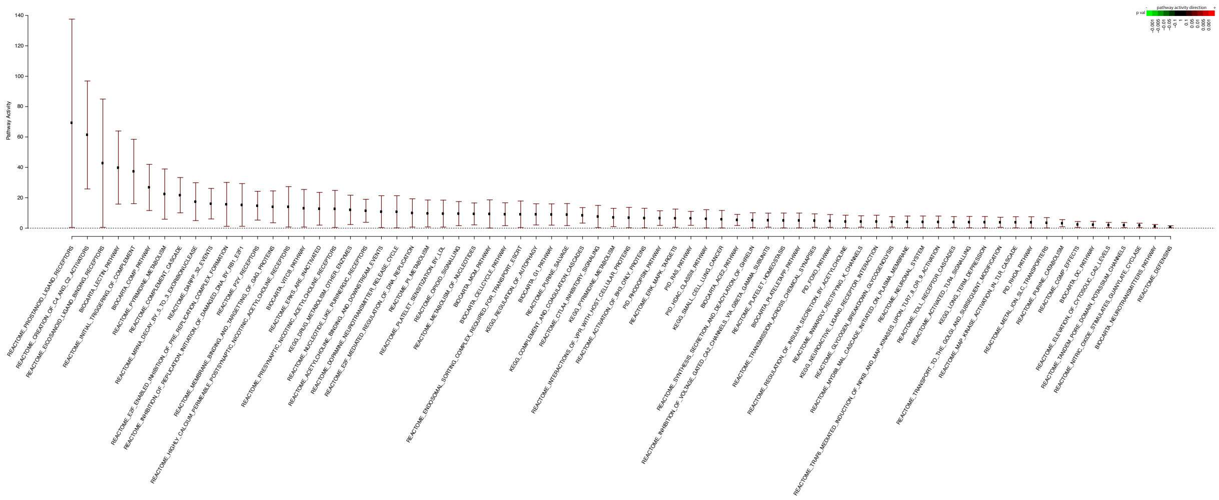

**B**

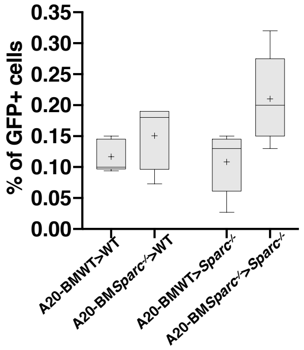
